## Supplementary figures and tables for "Structural analysis of SALL4 zinc-finger domain reveals a link between AT-rich DNA binding and Okihiro syndrome"

### Figure S1

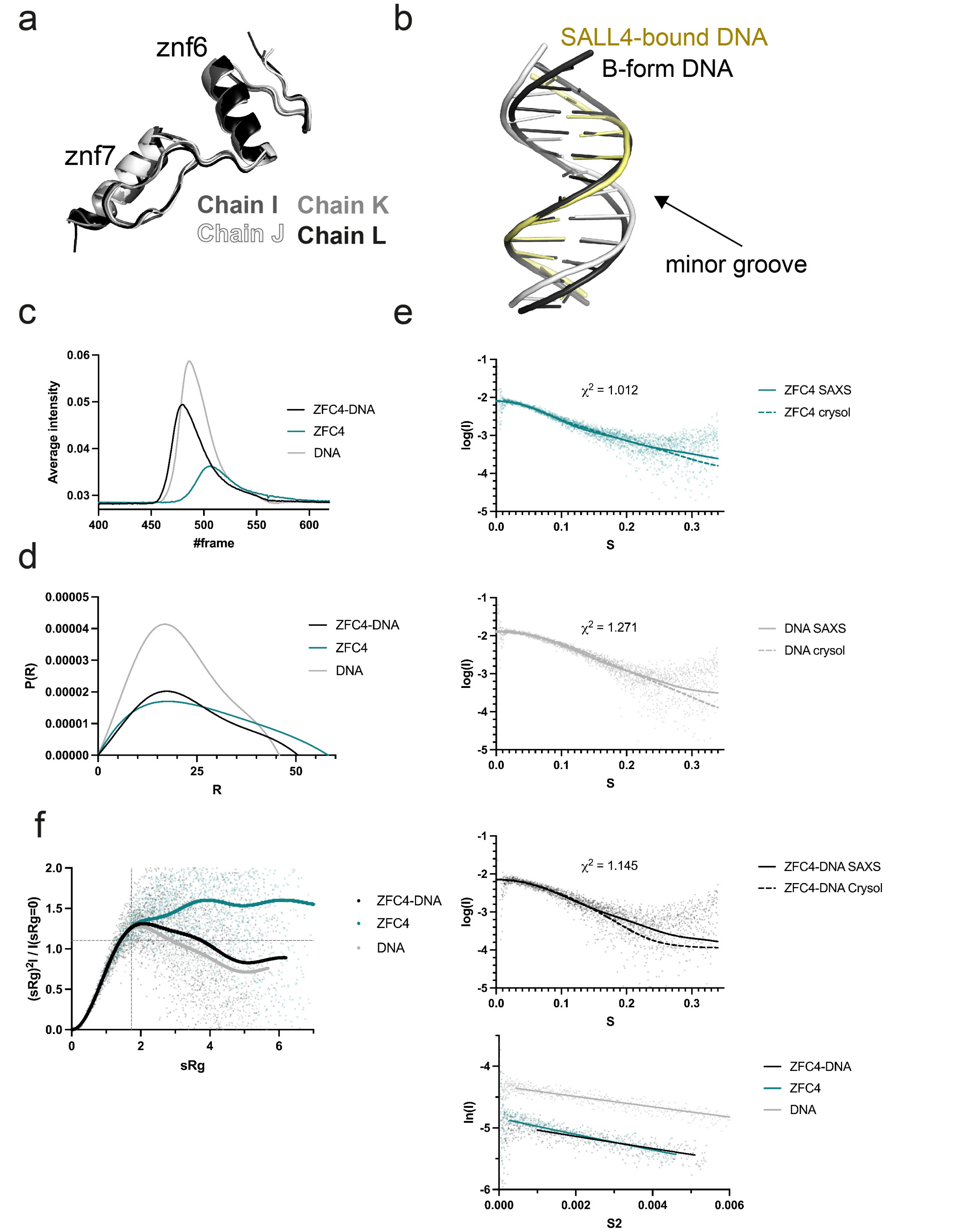


**Figure S1** (a) Superposition of all four ZFC4 chains found in the asymmetric unit of the crystal. (b) Superposition of refined AT-rich DNA (white and yellow) compared with ideal B-form (black). (c) SEC-SAXS profiles for SALL4 ZFC4 (teal), dsDNA (gray) and ZFC4-DNA complex (black). (d) Real space P(r) functions calculated for SAXS experiments in (c). (e) SAXS curves for samples from (c) showing comparison with curves calculated from all atom models of SALL4 ZFC4 (teal), DNA (grey) and the ZFC4-DNA complex (black). Guinier analysis for all samples are shown below. (f) Normalised Kratky analysis for SAXS samples.

### Figure S2

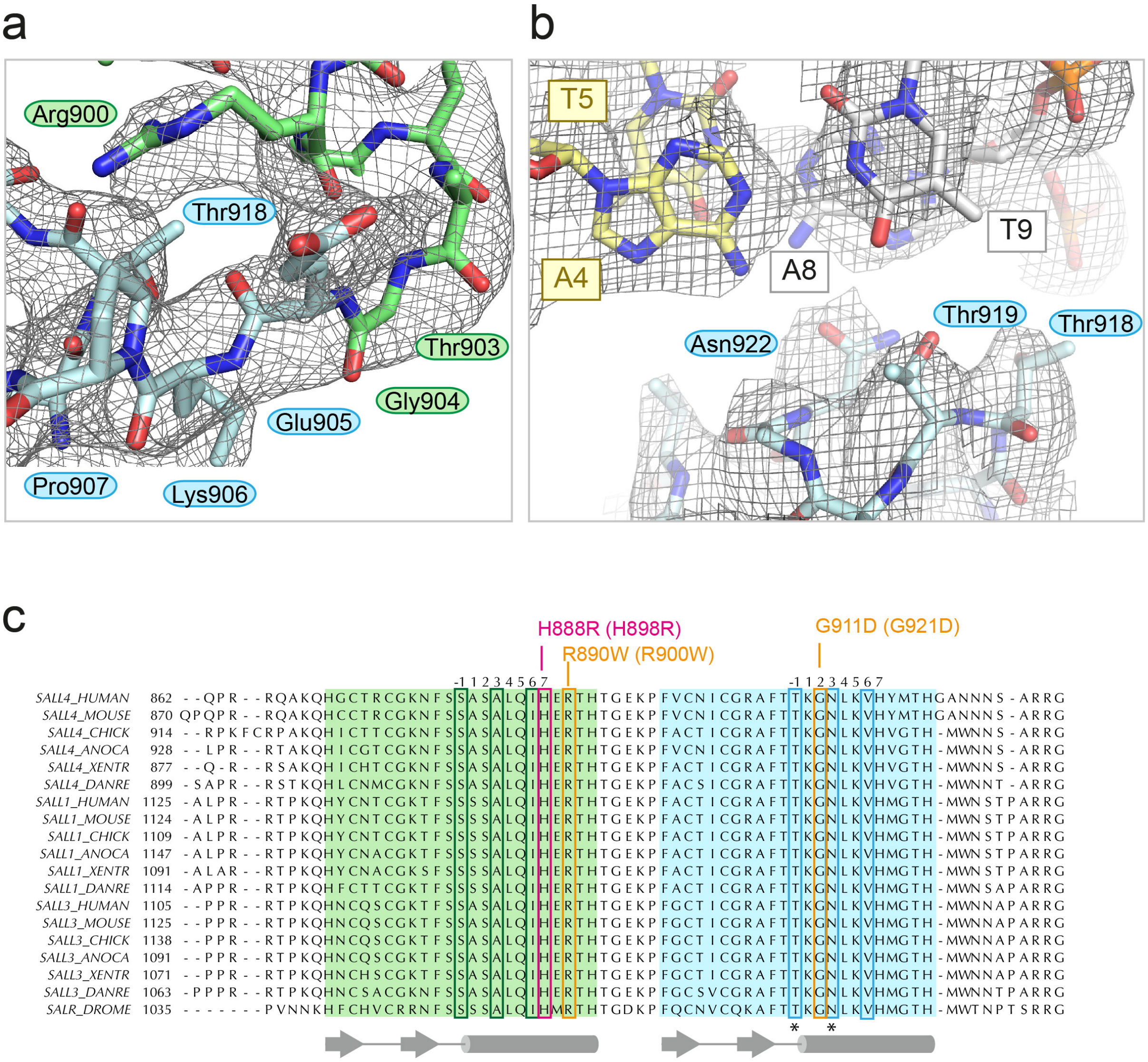


**Figure S2** (a) 2Fo-Fc map contoured at 1σ around the TGEKP sequence that connects Znf6 to Znf7. R900 of Znf6 hydrogen bonds with the backbone carbonyl group of T918. (b) Refined SALL4 ZFC4 chain L with palindromic DNA, showing the final 2Fo-Fc map contoured at 1σ. The view is of a plane through the A4(yellow)-T9(white) base pair. (c) Multiple protein sequence alignments of SALL4, SALL1, and SALL3 across vertebrates (*Homo sapiens* (HUMAN), *Mus musculus* (MOUSE), *Gallus gallus* (CHICK), *Anolis caroliensis* (ANOCA), *Xenopus tropicalis* (XENTR), *Danio rerio* (DANRE)), with *Drosophila melanogaster* Salr for reference. The sequences were aligned using MAFFT. Znf6 is highlighted green and Znf7 is highlighted blue. Reference sequence positions for each zinc finger are given above the alignment. DNA binding residues are boxed in green and blue. Patient missense mutations are boxed in pink and orange and annotated above with human and (mouse) numbering. Secondary structure elements are shown below the sequence. Asterisks indicate residues that have previously been shown to alter SALL4 function in mouse cells.

### Figure S3

a

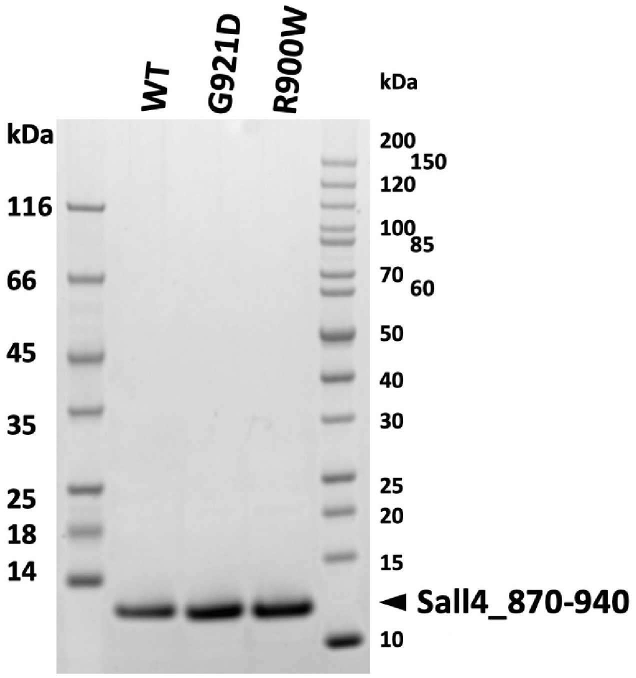


b

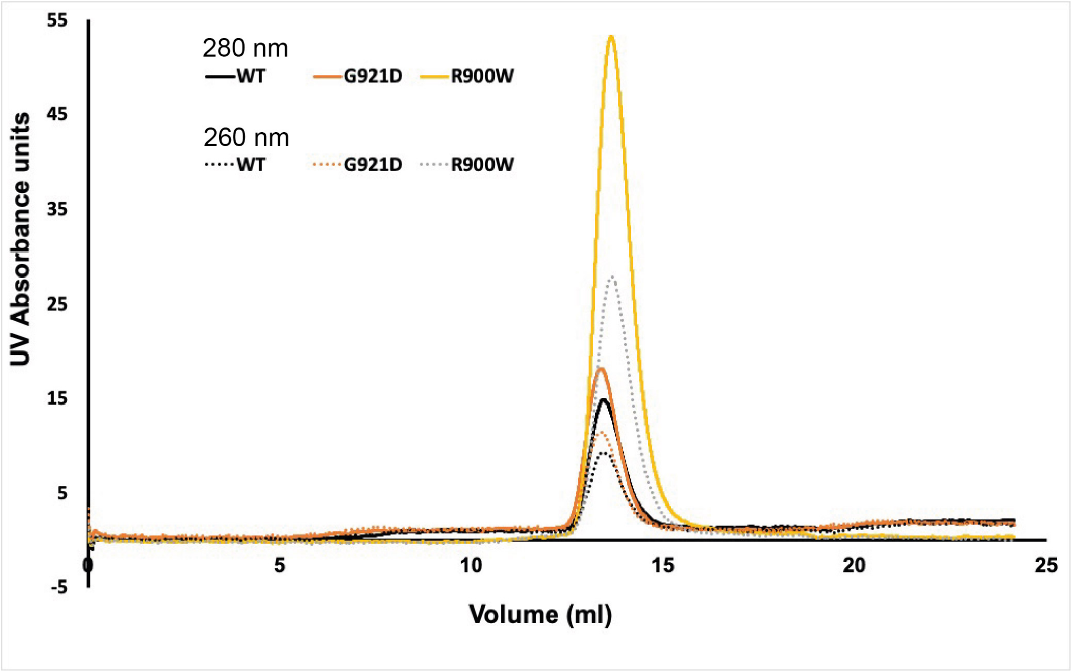


**Figure S3** (a) SDS-PAGE gel (4-20%) with wild-type and mutant SALL4 ZFC4 (residues 870-940). (b) Superposition of size exclusion chromatograms (from Superdex S75 column), showing elution profiles for wild-type (black), G921D (orange) and R900W (pale orange) SALL4 ZFC4.

Figure S4

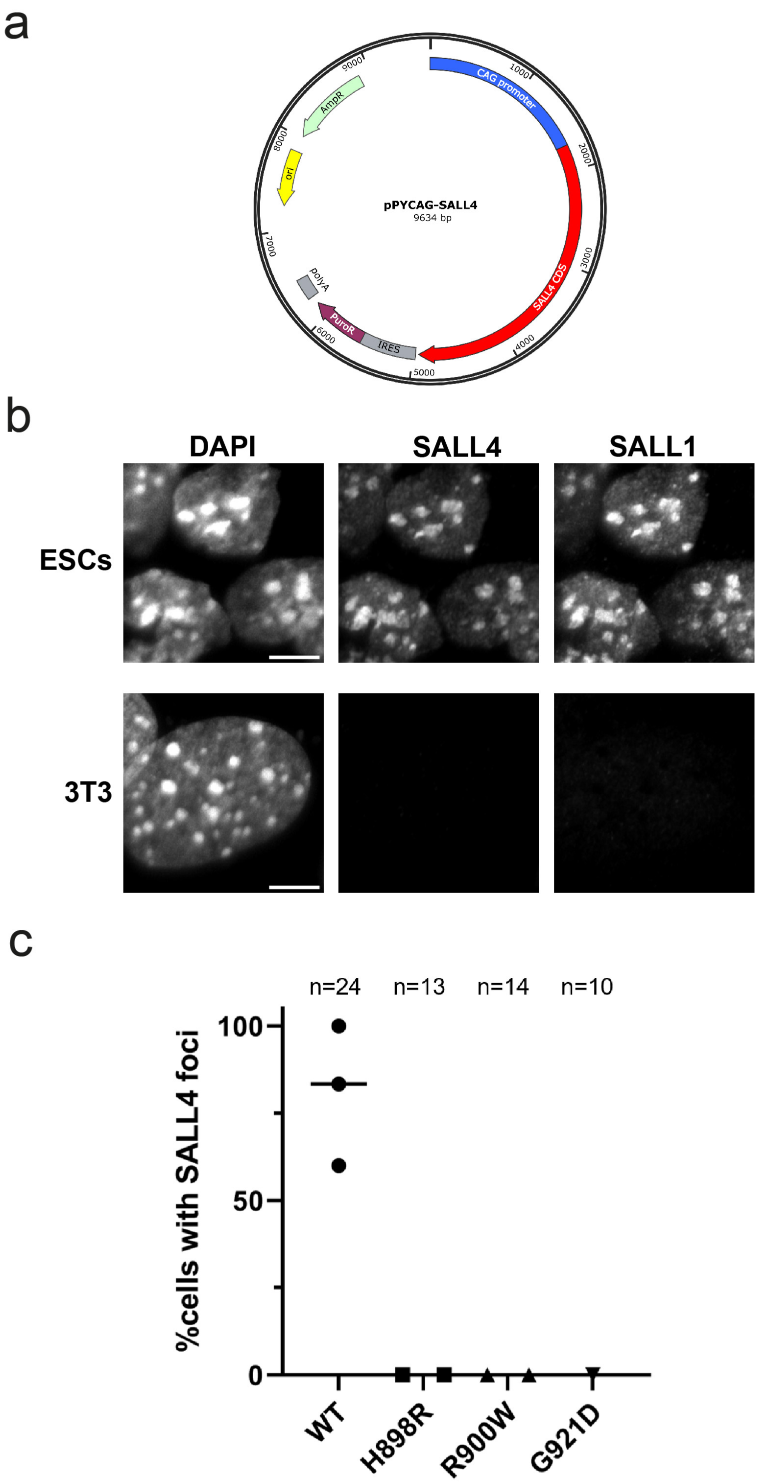


**Figure S4** (a) Plasmid used for expression of wild type and mutant SALL4 proteins in 3T3 cells. (b) DAPI staining and immunofluorescence for SALL4 and SALL1 in mouse embryonic stem cells (ESCs) and NIH 3T3 cells. Scale bars are 5µm. (c) Quantification of cells with SALL4 localisation to DAPI bright foci for wild type and mutant proteins. Total numbers of analysed cells are indicated for each construct.

### Figure S5

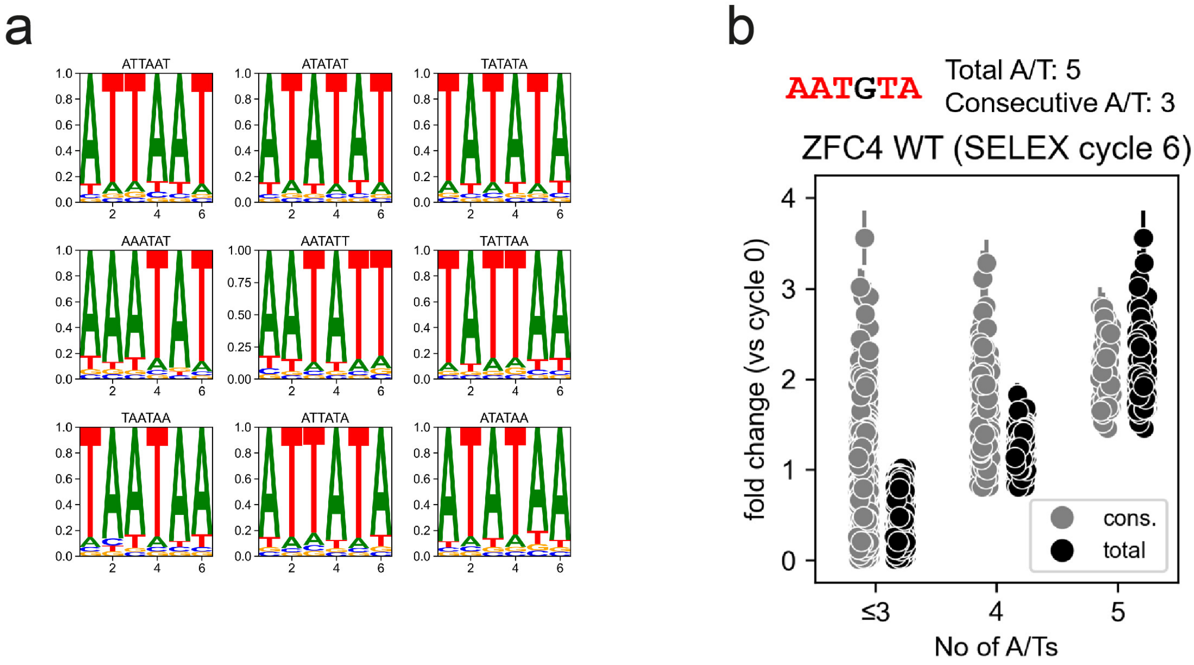


**Figure S5** (a) Position frequency matrix (PFM) motif logos of the most enriched 6-mer DNA motifs at cycle 6 of HT-SELEX with SALL4 ZFC4. (b) Relative enrichment (fold-change vs cycle 0) of 6-mer DNA motifs categorized by consecutive or total number of A/Ts (see example above the plot) at cycle 6 of HT-SELEX with SALL4 ZFC4 wild-type. Error bars indicate the variability (SD) in three independent replicate experiments.

**Table S1. Data collection and structural refinement statistics**

| Data collection | SALL4-pall01 complex |
|---|---|
| Source, wavelength | DLS I04, 1.28216 Å |
| Number of crystals | 1 |
| Space group | *P*1 |
| Cell parameters a, b, c (Å) | 39.03 66.11 77.94 |
| $\alpha, \beta, \gamma$ (°) | 73.04 76.43 76.14 |
| Resolution (Å) | 73.4-2.8 (3.3-2.78) |
| Multiplicity | 3.3 (3.1) |
| Average $I/\sigma(I)$ | 3.1 (1.9) |
| Completeness (ellipsoidal) (%) | 0.77 (0.31) |
| $R_{merge}$ | 0.45 (0.91) |
| $R_{meas}$ | 0.54 (1.10) |
| $R_{pim}$ | 0.29 (0.62) |
| $CC_{1/2}$ | 0.67 (0.89) |
| Wilson B factor (Å$^2$) | 45 |
| **Refinement** | |
| Unique reflections used for refinement | 6179 |
| $R_{work}/R_{free}$ (%) | 24.7 /25.4 |
| Average B factors (Å$^2$) | 72 Å$^2$ |
| R.m.s. deviations | |
| Bond lengths (Å) | 0.006 |
| Bond angles (°) | 0.83 |
| Ramachandran | |
| Favoured (%) | 95 |
| Additionally allowed (%) | 5 |
| Outliers (%) | 0 |

**Table S2 – SAXS experimental details and data parameters**

| (*a*) Sample details | ZFC4 | DNA | ZFC4-DNA |
|---|---|---|---|
| Organism | *M. musculus* | - | Combination of ZFC4 and DNA |
| Source (Catalogue No. or reference) | *E. coli* expressed | IDT DNA oligos | Combination of ZFC4 and DNA |
| Uniprot ID (residues in construct) + uncleaved tag | Q8BX22 870-940 +3C scar (GPDS) at N-terminus | 5'CATATTAATATC3' 3'GTATAATTATAG5' | Combination of ZFC4 and DNA |
| Extinction coefficient ε ($A_{280}$, 0.1%(w/v) | 0.210 | - | - |
| Molecular mass *M* from chemical composition (Da) | 8426 | 7446 | 15872 (assuming 1:1 binding) |
| SEC-SAXS column, s200 increase 3.2/200 | | | |
| Loading concentration (mg/ml) | 6.69 | 5.00 | 5.80 |
| Injection volume (µl) | 45 | 45 | 45 |
| Flow rate (ml/min) | 0.1 | 0.1 | 0.1 |
| Concentration measurement method | BCA assay | - | - |
| Solvent composition | 20 mM Tris pH 7.5 at 4°C, 200 mM NaCl | | |

| (*b*) SAS data collection parameters | |
|---|---|
| Instrument | Diamond Light Source Ltd Synchrotron. BL21 beamline, Eiger 4M detector (Dectris) |
| Wavelength (Å) | 0.9998 |
| Beam size at focal point (µm) | 34x40 |
| Sample-to-detector distance (m) | 4.014 |
| *q*-measurement range ($Å^{-1}$) | 0.0026 – 0.34 |
| Exposure time | 3 s |
| Sample temperature (ºC) | 22 |

| (*c*) Software employed for SAS data reduction, analysis and interpretation | |
|---|---|
| Sample – Solvent subtraction | Chromixs from ATSAS 3.0.5 (Manalastas-Cantos et al., 2021) |
| Calculation of ε from sequence | ProtParam (Gasteiger et al., 2005) |
| Basic analyses: Guinier, *P*(*r*), $V_P$ | PRIMUS from ATSAS 3.0.5 (Manalastas-Cantos et al., 2021) |
| Shape/bead modelling | DAMMIF from ATSAS 3.0.5 (Manalastas-Cantos et al., 2021) |
| | Crysol from PRIMUS in ATSAS 3.0.5 (Manalastas-Cantos et al., 2021) |
| Modelling of missing sequence | COOT (Emsley and Cowtan, 2004) |
| Molecular graphics | PyMOL 2.4.1 and ChimeraX 1.3 |

| (*d*) Structural parameters | ZFC4 | DNA | ZFC4-DNA |
|---|---|---|---|
| Guinier Analysis | | | |
| *I*(0) ($cm^{-1}$) | 0.082 ± 0.00005 | 0.014 ± 0.000069 | 0.0075 ± 0.000077 |
| $R_g$ (Å) | 20.77 ± 0.24 | 16.72 ± 0.17 | 18.23 ± 0.36 |
| q range ($Å^{-1}$) | 0.0026 – 0.340 | 0.0026 – 0.340 | 0.0026 – 0.340 |
| Quality of Fit $\chi^2$ value (p-value) | 1.24 (0.00692) | 1.08 (0.198) | 0.857 (0.953) |

| | | | |
|---|---|---|---|
| $P(r)$ analysis | | | |
| $I(0)$ ($cm^{-1}$) | 0.0079 ± 0.0000394 | 0.0133 ± 0.0000536 | 0.00719 ± 0.0000608 |
| $R_g$ (Å) | 20.08 ± 0.0819 | 16.02 ± 0.0634 | 17.48 ± 0.124 |
| $d_{max}$ (Å) | 58.22 | 45.82 | 50.56 |
| $q$-range (Å$^{-1}$) | 0.00412 – 0.340 | 0.00487 – 0.340 | 0.00675 – 0.340 |
| Quality of Fit $\chi^2$ value (p-value) | 1.01 (0.351) | 1.03 (0.124) | 1.01 (0.322) |
| Porod volume (Å$^{-3}$) | 8564.57 | 12052.90 | 14565.90 |

| (*e*) Shape modelling results | ZFC4 | DNA | ZFC4-DNA |
|---|---|---|---|
| DAMMIF (default parameters, 10 repetitions), averaged with DAMAVER and refined with DAMMIN | | | |
| $q$-range for fitting | 0.00412 – 0.340 | 0.00487 – 0.340 | 0.00675 – 0.340 |
| Symmetry/anisotropy assumptions | P1, none | P1, none | P1, none |
| NSD (standard deviation) | 0.753 (0.044) | 0.752 (0.046) | 0.754 (0.055) |
| $\chi^2$ value | 0.9915 | 1.032 | 1.013 |
| Constant subtraction procedure | Skipped | 0.0001656 | 0.0001076 |
| Model resolution (from SASRES) (Å) | 20 | 17 | 20 |

| (*f*) Atomistic modelling | ZFC4 | DNA | ZFC4-DNA |
|---|---|---|---|
| CRYSOL (with default parameters) | | | |
| $q$-range for fitting | | | |
| $\chi^2$ value (p-value) | 1.012 (0.333) | 1.272 ($1.58 \times 10^{-19}$) | 1.145 ($3.35 \times 10^{-7}$) |
| Predicted Rg (Å) | 21.56 | 14.28 | 16.88 |
| Vol (Å), Ra (Å), Dro (e Å$^{-3}$) | 10008, 1.400, 0.068 | 7717, 1.400, 0.075 | 17725, 1.800, 0.000 |
| (*g*) Data and model deposition IDs | pending | pending | pending |


Manalastas-Cantos, K., Konarev, P.V., Hajizadeh, N.R., Kikhney, A.G., Petoukhov, M.V., Molodenskiy, D.S., Panjkovich, A., Mertens, H.D.T., Gruzinov, A., Borges, C., Jeffries, C.M., Svergun, D.I., Franke, D. (2021)
ATSAS 3.0: expanded functionality and new tools for small-angle scattering data analysis *J. Appl. Cryst.* 54, 343-355 DOI

Gasteiger E., Hoogland C., Gattiker A., Duvaud S., Wilkins M.R., Appel R.D., Bairoch A.;
*Protein Identification and Analysis Tools on the ExPASy Server;*
(In) John M. Walker (ed): The Proteomics Protocols Handbook, Humana Press (2005).
pp. 571-607

Emsley, P. and K. Cowtan (2004). "Coot: model-building tools for molecular graphics." Acta Crystallogr D Biol Crystallogr 60(Pt 12 Pt 1): 2126-2132.

**Table S3 Antibody dilutions for immunofluorescence**

| Target | Application | Product reference | Working dilution |
|---|---|---|---|
| SALL4 | Immunofluorescence | Abcam cat. ab29112 | 1:200 |
| SALL4 | Immunofluorescence | Santa Cruz cat. sc-101147 | 1:50 |
| SALL1 | Immunofluorescence | Abcam cat. ab41974 | 1:200 |
